# Incomplete activation of developmentally required genes *Alyref1* and *Gabpb1* leads to preimplantation arrest in cloned mouse embryos

**DOI:** 10.1101/2022.04.14.488417

**Authors:** Shunya Ihashi, Mizuto Hamanaka, Masaya Kaji, Miki Mori, Yuma Imasato, Misaki Nakamura, Masayuki Anzai, Kazuya Matsumoto, Masahito Ikawa, Kei Miyamoto

## Abstract

Differentiated cell nuclei can be reprogrammed after nuclear transfer (NT) to oocytes and the produced NT embryos can give rise to cloned animals. However, development of NT embryos is often hampered by recurrent reprogramming failures, including the incomplete activation of developmental genes, yet specific genes responsible for the arrest of NT embryos are not well understood. Here, we searched for developmentally important genes among the reprogramming-resistant H3K9me3-repressed genes, and identified *Alyref* and *Gabpb1* by siRNA screening. Gene knockout of *Alyref* and *Gabpb1* by the CRISPR/Cas9 system resulted in early developmental arrest in mice. Single embryo RNA-seq revealed that *Alyref* is needed for the formation of inner cell mass. The supplement of *Alyref* and *Gabpb1* by mRNA injection supported efficient preimplantation development of cloned embryos. Thus, our study shows that the H3K9me3-repressed genes contain developmentally required genes and the incomplete activation of such genes results in preimplantation arrest of cloned embryos.

## INTRODUCTION

After fertilization, germ cells acquire totipotency, the ability to give rise to all cells that build the conceptus. On the other hand, a somatic cell nucleus can be reprogrammed to be totipotent after transplanting it into an enucleated oocyte. This technique is called somatic cell nuclear transfer (SCNT). Using SCNT, the successful cloning of an animal was first reported in *Xenopus laevis* (Gurdon, 1962), followed by various mammalian species such as sheep, mice, and cattle (Cibelli et al., 1998; Wakayama et al., 1998; Wilmut et al., 1997). However, the efficiency of obtaining cloned animals by conventional SCNT methods is low, and a myriad of defects have been reported during both pre- and post-implantation development (Loi et al., 2016; Ogura et al., 2013). The low efficiency of SCNT is one of the main factors that impede the prevalence of this technology, even though SCNT potentially offers an invaluable opportunity for basic biology, regenerative medicine, and the propagation of endangered animals.

The molecular basis for the low developmental potential of SCNT embryos has been investigated. It is now generally recognized that the abnormal state of histone modifications in SCNT embryos is a major cause of aberrant gene expression and low developmental ability. Facilitation of histone acetylation by the addition of Trichostatin A (TSA), a small molecule compound that inhibits histone deacetylases, into embryo culture medium has been widely applied for enhancing development of SCNT embryos (Kishigami et al., 2006; Rybouchkin et al., 2006). It has also been reported that histone H3 lysine 9 trimethylation (H3K9me3) accumulated in somatic genomes persists after SCNT and prevents zygotic genome activation (ZGA) of a subset of genes (Matoba et al., 2014). Inhibitory effects of H3K9me3 on nuclear reprogramming in SCNT embryos have been confirmed in several species (Chung et al., 2015; Liu et al., 2018). Furthermore, the persisting H3K9me3 mark prevents nuclear reprogramming in induced pluripotent stem (iPS) cells (Soufi et al., 2012), demonstrating H3K9me3 as an universal barrier to reprogramming. In other words, H3K9me3 serves as a solid mechanism to repress lineage-inappropriate genes in somatic cell genomes (Becker et al., 2016), and it would be important to experimentally test if genes required for accomplishing reprogramming are indeed included among H3K9me3-repressed genes.

In our previous study, we found that when mouse SCNT embryos are treated with TSA and then cultured in vitamin C (VC)-supplemented medium, 83.6% of SCNT embryos develop to the blastocyst stage and 15.2% of SCNT embryos develop into offspring (Miyamoto et al., 2017), representing one of the most efficient development of cloned embryos (Ogura, 2020). This treatment with VC lowers the level of H3K9me3 and significantly alters transcriptomes in SCNT embryos (Miyamoto et al., 2017). It is therefore plausible that reprogramming-resistant genes regulated by H3K9me3 are upregulated by the TSA and VC treatment, which might have eventually resulted in the enhanced development of SCNT embryos. In this paper, we found that among previously identified reprogramming-resistant genes (Matoba et al., 2014), 16 genes were upregulated in SCNT embryos under TSA and VC-treated conditions. We further performed siRNA screening to identify developmentally important genes, and *Alyref* and *Gabpb1* were identified. Alyref is an mRNA-binding adaptor protein involved in nuclear export of mRNA (Rodrigues et al., 2001; Zhou et al., 2000). Alyref interacts with lws1 to mediate mRNA transport (Yoh et al., 2007), and it has been reported that the knockdown of lws1 reduces mRNA transport and arrests embryonic development at 8-16 cell stage (Oqani et al., 2019). *Gabpb1* encodes Gabp-β, one of the subunits of GA-binding protein (GABP) that acts as a transcriptional activator (Rosmarin et al., 2004). GABP is known to activate *Yap* gene, which is necessary for preimplantation development (Wu and Guan, 2021), and the knockdown of GABP has been reported to decrease YAP and inhibit progression to G1/S stage, eventually leading to increased cell death (Wu et al., 2013). However, roles of *Alyref* and *Gabpb1* in preimplantation development remain unclear. Here, we showed that knockout of *Alyref* and *Gabpb1* caused early embryonic arrest in mouse fertilized embryos. Furthermore, expression of Alyref and Gabpb1 was repressed in SCNT embryos at the protein level, and rescued expression of Alyref and Gabpb1 by mRNA injection enhanced preimplantation development of SCNT embryos. Therefore, *Alyref* and *Gabpb1*, genes necessary for preimplantation development, are normally repressed in SCNT embryos, and the incomplete activation of *Alyref* and *Gabpb1* at least partially explains the low developmental capacity of SCNT embryos.

## RESULTS

### Identification of developmentally important genes that are downregulated in SCNT embryos

We have previously shown that development of SCNT embryos is greatly enhanced by the combinational treatment of TSA and VC (Miyamoto et al., 2017). We hypothesized that these epigenetic modifiers allowed activation of key genes for embryonic development, which were otherwise repressed in SCNT embryos. We therefore compared differentially expressed genes (DEGs) found by the TSA+VC treatment (control SCNT vs SCNT with TSA and VC) (Miyamoto et al., 2017) to those found between *in vitro* fertilized (IVF) embryos and SCNT embryos (a group of 301 genes that failed to be activated in SCNT embryos) (Matoba et al., 2014). Among the DEGs, 16 genes were shared (Figures 1A and 1B). These genes are normally repressed in SCNT embryos, but are upregulated when the development of cloned embryos is greatly enhanced by the TSA+VC treatment. We then searched for developmentally important genes among the list by knocking down their expression in IVF embryos with siRNA injection and specific siRNA sets for 15 genes were designed. The siRNA screening identified that knockdown of *Alyref* and *Gabpb1* significantly impaired the development of IVF embryos to the blastocyst stage (Figure 1C). Successful knockdown of both genes was shown by reverse transcription-quantitative polymerase chain reaction (RT-qPCR) (Figure S1A). We also observed a significant decrease in preimplantation development when the mixture of siRNAs that target transcriptional activators, including *Gabpb1*, as shown in Figure 1B (red colors) were injected (Figure 1C). Inhibition of *Alyref* or *Gabpb1* by siRNA injection resulted in developmental arrest at the morula stage (Figures 1D and S1B); especially almost all embryos did not develop to the blastocyst stage in *Alyref* siRNA-injected embryos (Figure S1B). These results suggest that *Alyref* or *Gabpb1* genes, repressed in SCNT embryos, are important for the progression to the blastocyst stage.

**Figure 1.**
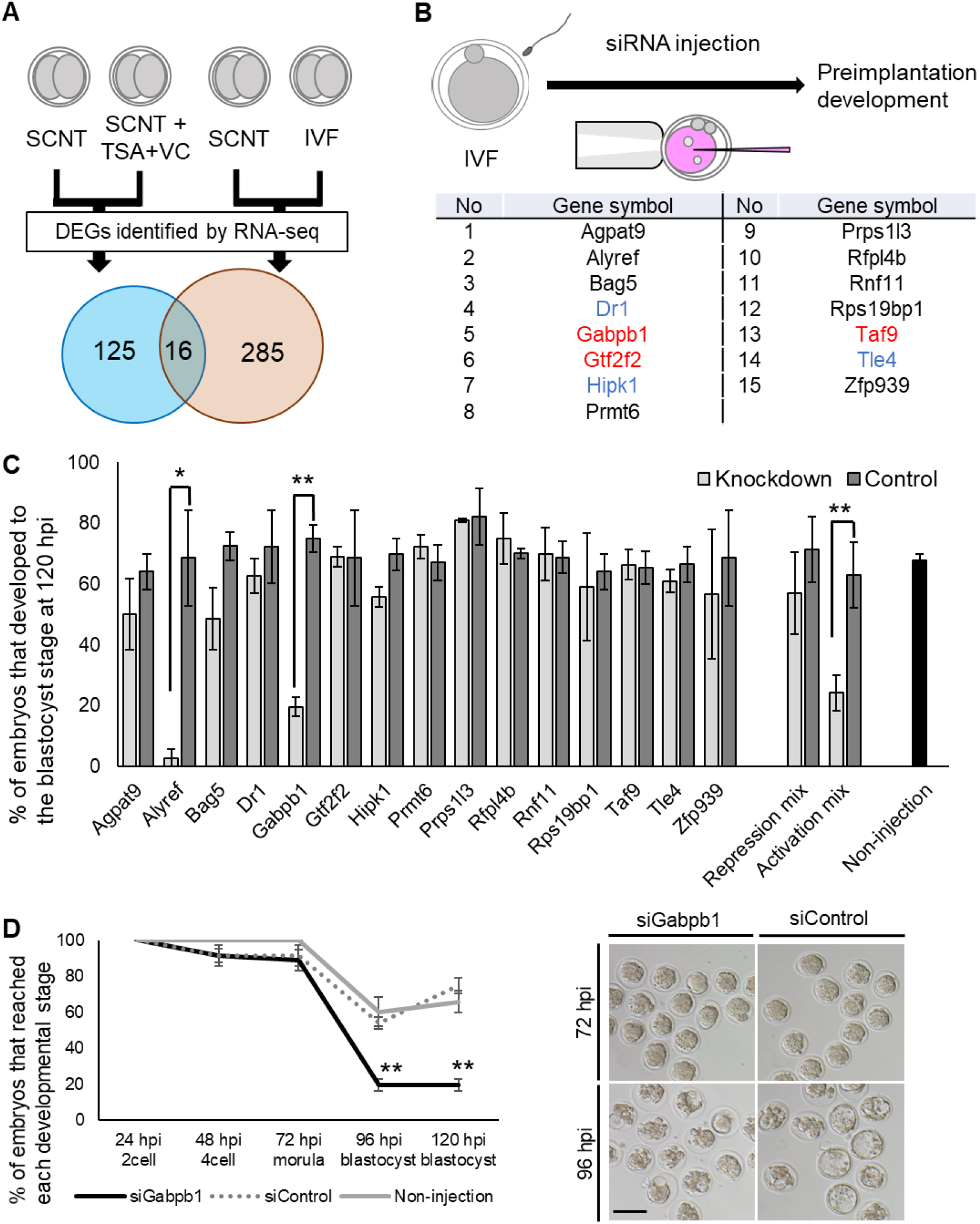
Screening for candidate genes key for development of mouse SCNT embryos. (A) A strategy for identifying genes responsible for reprogramming in SCNT embryos and those for development in normal IVF embryos. (B) An experimental scheme for siRNA screening to find genes important for development of fertilized embryos and the list of candidate genes. Genes related to gene activation are shown in red, while those related to gene repression are in blue. (C) Development of embryos injected with the listed siRNAs. Control represents embryos injected with control siRNA. Repression mix includes genes marked with blue in Figure 1B, while activation mix includes those with red. Statistical significance was calculated by F- and T-test. Three to eight independent experiments were performed. Error bars are SE. **p < 0.01, *p < 0.05. (D) Preimplantation development of embryos injected with siRNA against *Gabpb1*, control siRNA, and non-injected embryos. Representative images of the injected embryos are shown in the right panel. Statistical significance was calculated by chi-square test. The numbers of embryos used for experiments are as follows; siGabpb1: 55, siControl: 48, and Non injection: 21. Four independent experiments were performed. Error bars are SE. Scale bar, 100 μm. **p < 0.01.

### Abnormal expression of Alyref and Gabpb1 in SCNT embryos

We next examined if abnormal expression of Alyref or Gabpb1 is indeed observed in SCNT embryos. Firstly, the protein localization of Alyref and Gabpb1 was examined during the preimplantation development. Alyref and Gabpb1 were localized in pronuclei of mouse IVF zygotes (Figure 2A), suggesting maternal accumulation of Alyref and Gabpb1 protein. Nuclear localization was seen throughout the preimplantation development. To note, dot-like localization of Alyref was observed in nuclei of IVF embryos at the 2-cell stage onwards (Figures 2A and S2A). Gabpb1 protein was accumulated around nucleoli at the 4-cell stage onwards (Figure 2B). Secondly, transcriptional expression of *Alyref* or *Gabpb1* in SCNT embryos was examined using the published RNA-seq dataset (Matoba et al., 2014; Miyamoto et al., 2017). Both *Alyref* and *Gabpb1* were repressed in SCNT embryos compared to IVF embryos at the 2-cell stage and this repression was rescued by the overexpression of Kdm4d (Figures S2B and S2C), which removes excess H3K9me3 observed in SCNT embryos (Matoba et al., 2014). Upregulation of *Alyref* and *Gabpb1* was also observed in SCNT embryos treated with TSA and VC (Figures S2B and S2C), which also lowers the level of H3K9me3 (Miyamoto et al., 2017). These results suggest that Alyref and Gabpb1 are repressed at the transcription level in SCNT embryos at the time of ZGA. Lastly, protein expression of Alyref and Gabpb1 in SCNT embryos was examined. Both Alyref and Gabpb1 were significantly downregulated in SCNT embryos especially at the morula stage when compared to IVF embryos (Figures 2C and 2D). These results suggest that abnormal downregulation of Alyref and Gabpb1 is observed in SCNT embryos both at the transcript and protein levels.

**Figure 2.**
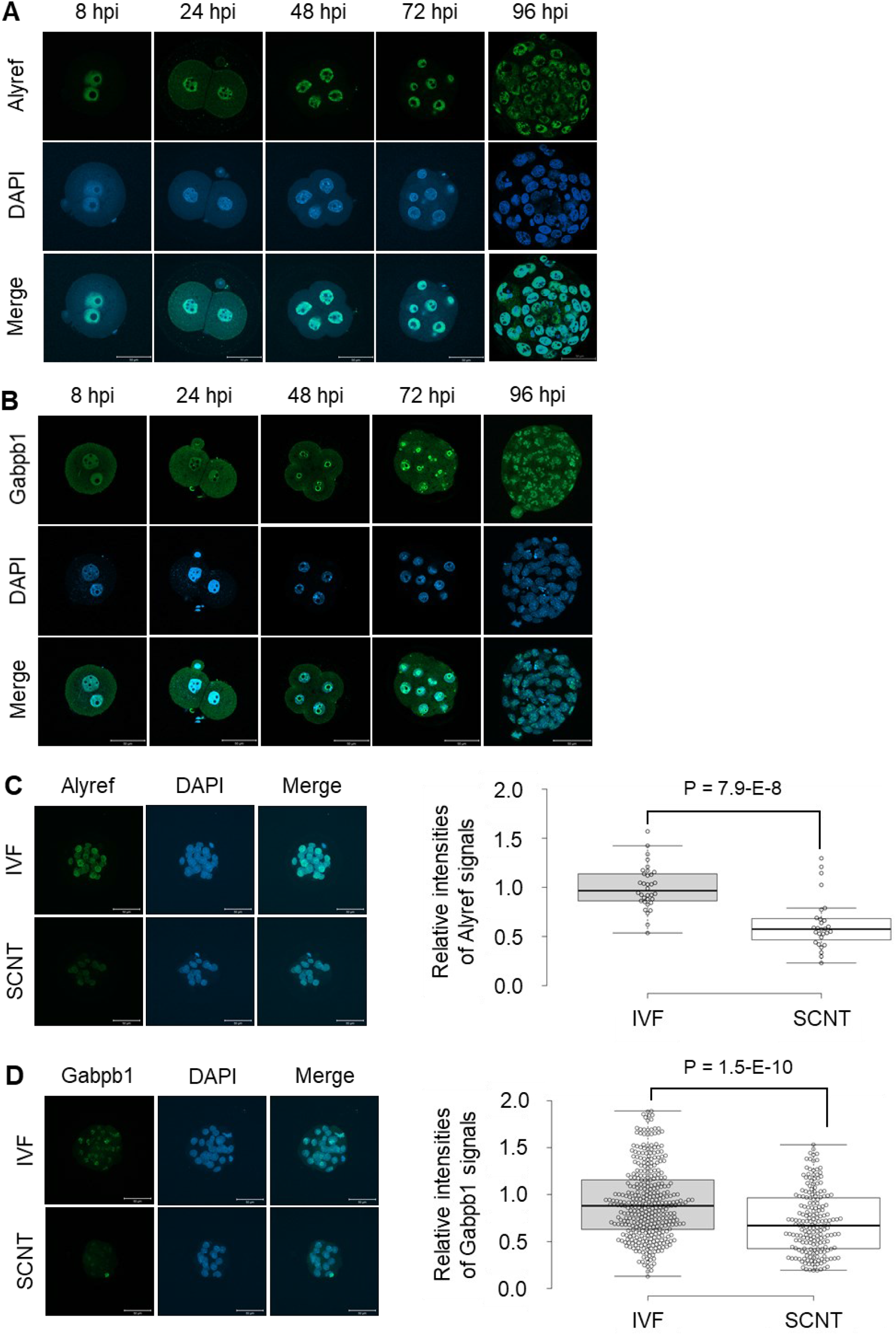
Localization of Alyref and Gabpb1 proteins in fertilized and cloned embryos. (A) Immunostaining of Alyref in fertilized embryos during preimplantation development. DNA was stained with DAPI. Merge represents the merged photos. n = 6∼19 Three independent experiments were performed. Scale bars, 50 μm. (B) Immunostaining of Gabpb1 in fertilized embryos during preimplantation development. n = 36∼63. Four to Six independent experiments were performed. Scale bars, 50 μm. (C) Immunostaining of Alyref at the morula stage (left panel) and the box plot indicates signal intensities of Alyref in IVF and SCNT embryos. The numbers of embryos used for experiments are as follows: SCNT: 28, IVF: 34. Five independent experiments were performed. Two-sided T test was used. Scale bars, 50 μm. (D) Immunostaining of Gabpb1 at the morula stage (left panel) and the box plot indicates signal intensities of Gabpb1 in IVF and SCNT embryos. The numbers of embryos used for experiments are as follows: SCNT: 23, IVF: 40. Five independent experiments were performed. Two-sided T test was used. Scale bars, 50 μm.

### *Alyref* gene is essential for preimplantation development

To clarify the requirement of *Alyref* for embryonic development, we generated knockout mice by the CRISPR/Cas9 gene-editing method. The coding region of *Alyref* gene was removed by introducing gRNAs and Cas9 protein to zygotes as reported before (Mashiko et al., 2013) (Figure 3A). The resulting hetero mice (Alyref^+/-^) were identified by PCR-based genotyping (Figure S3A). When Alyref^+/-^ mice were mated with wild type mice, viable offspring was obtained (Figure 3B). However, the number of offspring was significantly reduced (Figure S3B) and no knockout mice (Alyref^-/-^) were obtained after crossing Alyref^+/-^ x Alyref^+/-^ (Figure 3B), suggesting the embryonic lethal phenotype of *Alyref* knockout mice. We then performed IVF using sperm and oocytes obtained from Alyref^+/-^ mice in order to reveal when development is arrested. The significant decrease in percentages of embryos that reach the blastocyst stage was observed (Figures 3C and 3D). Immunofluorescent analyses revealed that all blastocyst embryos obtained by Alyref^+/-^ x Alyref^+/-^ showed Alyref signals, while some of the morula embryos lacked Alyref expression (Figure S3C). These results demonstrate that *Alyref* is required for mouse preimplantation development and knockout of *Alyref* causes developmental arrest at the morula stage.

**Figure 3.**
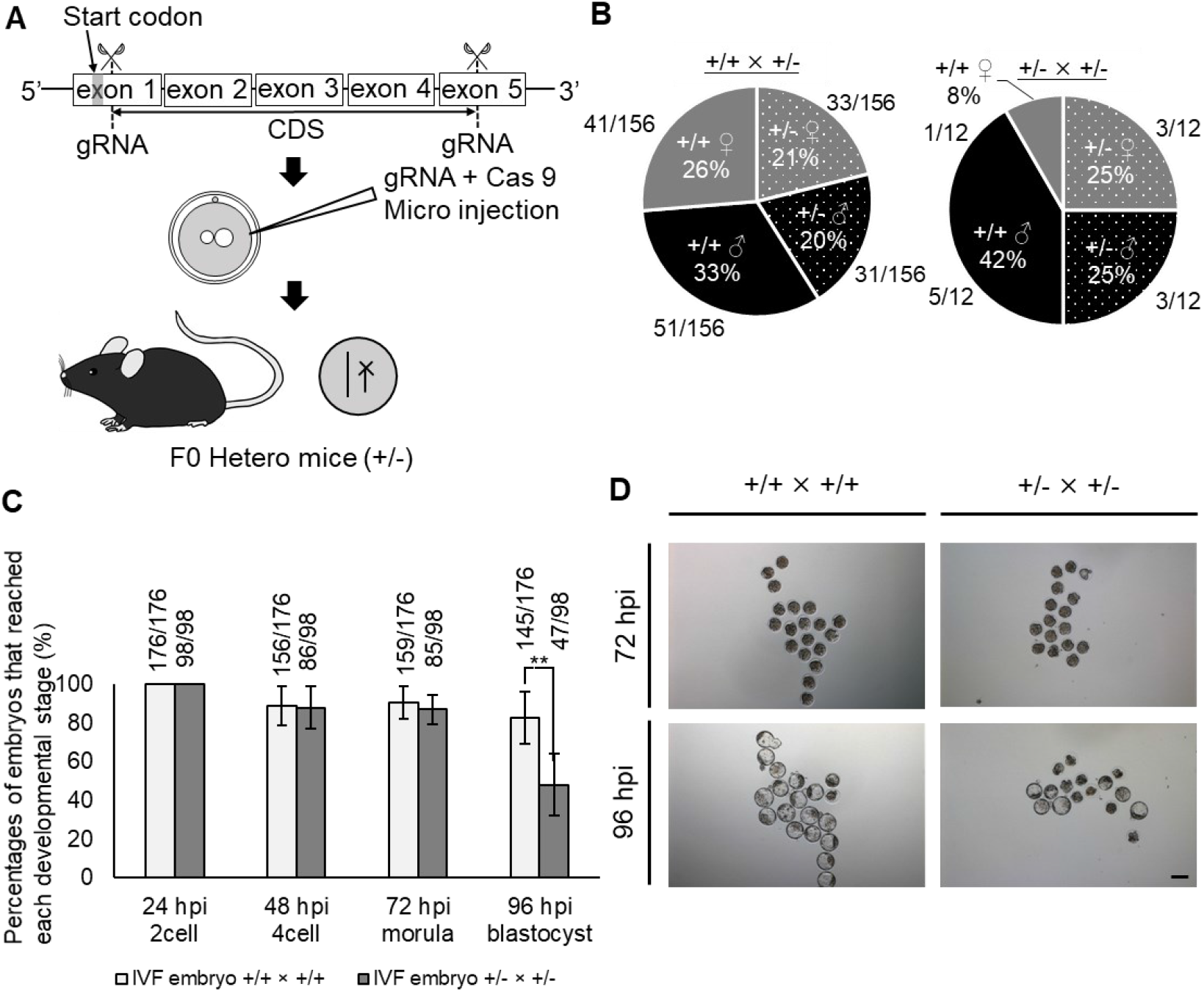
*Alyref* is required for mouse preimplantation development. (A) A scheme for CRISPR/Cas9-mediated knockout of *Alyref* gene. Two gRNAs were introduced with Cas9 protein to mouse zygotes in order to induce the large deletion of the coding region of *Alyref*. (B) Pie charts represent percentages of genotypes and sex of offspring after each mating (Alyref^+/+^ x Alyref^+/-^ [left] and Alyref^+/-^ x Alyref^+/-^ [right]). The numbers of offspring are also shown. (C) Preimplantation development of wild type embryos (Alyref^+/+^ x Alyref^+/+^) and embryos generated by mating heterozygous mice (Alyref^+/-^ x Alyref^+/-^). The numbers of embryos that reached each developmental stage are indicated above the bars. Error bars are SE. Statistical significance was calculated by chi-square test (**p < 0.01). Six independent experiments were performed. (D) Representative images of embryos. Control Alyref^+/+^ x Alyref^+/+^ embryos are shown in left panels, while Alyref^+/-^ x Alyref^+/-^ embryos are shown in right. Scale bar, 100 μm.

### *Gabpb1* gene is essential for embryonic development

To test the requirement of *Gabpb1* for embryonic development, we generated knockout mice by the CRISPR/Cas9 method. gRNA that targets exon 3 of *Gabpb1* was designed and introduced to zygotes with Cas9 protein (Figure 4A). The resulting hetero mice (Gabpb1^+/-^) were identified by DNA sequencing (Figure S4A) and the frame shift mutation was confirmed in the knockout allele (Figure 4B), which resulted in a truncated protein. When Gabpb1^+/-^ mice were mated with wild type mice, viable offspring was obtained (Figures 4C and 4D). However, when Gabpb1^+/-^ mice were mated with Gabpb1^+/-^, the number of offspring was significantly reduced (Figure 4D) and no knockout mice (Gabpb1^-/-^) were obtained (Figure 4C). We then carried out IVF using sperm and oocytes obtained from Gabpb1^+/-^ mice. The significant decrease in percentages of embryos that reach the morula and blastocyst stage was observed (Figures 4E and S4B).

**Figure 4.**
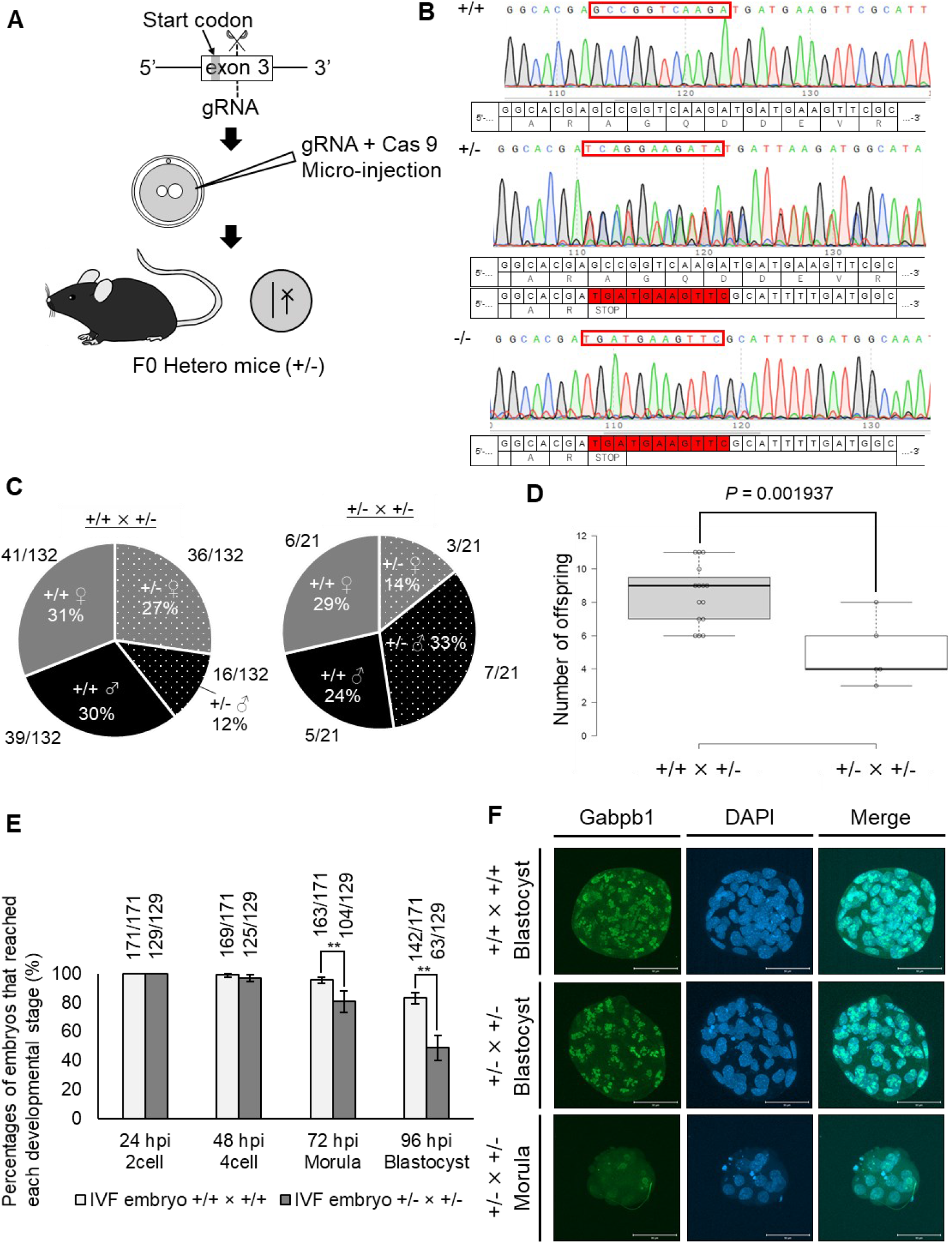
*Gabpb1* is required for mouse embryonic development. (A) A scheme for CRISPR/Cas9-mediated knockout of *Gabpb1* gene. gRNA that targets exon 3 of *Gabpb1* was introduced with Cas9 protein to mouse zygotes in order to induce the frame shift mutation of *Gabpb1*. (B) DNA sequencing analyses show the frame shift mutation in the knockout allele of *Gabpb1*. The deleted genomic region is highlighted in red color. (C) Pie charts represent percentages of genotypes and sex of offspring after each mating (Gabpb1^+/+^ x Gabpb1^+/-^ [left] and Gabpb1^+/-^ x Gabpb1^+/-^ [right]). The numbers of offspring are also shown. (D) The number of offspring is shown after each mating. Statistical significance was calculated by two-sided T test. (E) Preimplantation development of wild type IVF embryos (Gabpb1^+/+^ x Gabpb1^+/+^) and IVF embryos generated by using heterozygous mice (Gabpb1^+/-^ x Gabpb1^+/-^). The numbers of embryos that reached each developmental stage are indicated above the bars. Statistical significance was calculated by chi-square test (**p < 0.01). Error bars represent SE. Six independent experiments were performed. (F) Immunostaining of Gabpb1 in IVF embryos derived from different combinations of crossing. DNA was stained with DAPI. The numbers of embryos used for experiments are as follows: (Gabpb1^+/+^ x Gabpb1^+/+^): 102, and (Gabpb1^+/-^ x Gabpb1^+/-^): 82. Three independent experiments were performed. Scale bars, 50 μm.

Nevertheless, a certain number of embryos reached blastocysts after crossing Gabpb1^+/-^ x Gabpb1^+/-^, and we have therefore performed genotyping of the generated embryos. Many of Gabpb1^-/-^ embryos (75%) were arrested at the morula stage or degenerated, while 61% of Gabpb1^+/-^ embryos developed to the blastocyst state (Figure S4C). The lack of Gabpb1^-/-^ expression in the arrested morula embryos was also confirmed by immunofluorescent analyses (Figure 4F). These results demonstrate that Gabpb1 is required for mouse embryonic development.

### Developmentally important pathways are disturbed after knocking out *Alyref* or *Gabpb1*

To gain mechanistic insight into the role of *Alyref* and *Gabpb1*, we performed RNA sequencing (RNA-seq) analysis. Based on the IVF experiments using sperm and oocytes of heterozygous mice, most of knockout embryos were arrested at the morula stage at 96 hpi (Figures 3 and 4). Therefore, embryos remained as morulae at 96 hpi were collected and subjected to single embryo RNA-seq together with control blastocyst embryos (Figure 5A). The large deletion of *Alyref* was confirmed in two morula embryos (Figure 5B; KOA-M1 and KOA-M2) since no sequence reads were detected at the deleted region. Similarly, the frameshift mutation of *Gabpb1* was also confirmed in three morula embryos (Figure 5C; KOG-M1-M3). Off target deletion by CRISPR/Cas9 was not observed in *Alyref2* and *Gabpb2*, family genes of *Alyref* and *Gabpb1*, respectively (Figures S5A and S5B). PCA and hierarchical clustering analyses indicated that knockout morulae showed distinct transcriptomes from wild type morulae and were well separated from those of blastocyst embryos (Figures 5D, 5E, S5C and S5D). These results suggest that knockout of *Alyref* and *Gabpb1* causes abnormal gene expression in morulae, which may prevent the progression of knockout embryos to the blastocyst stage.

**Figure 5.**
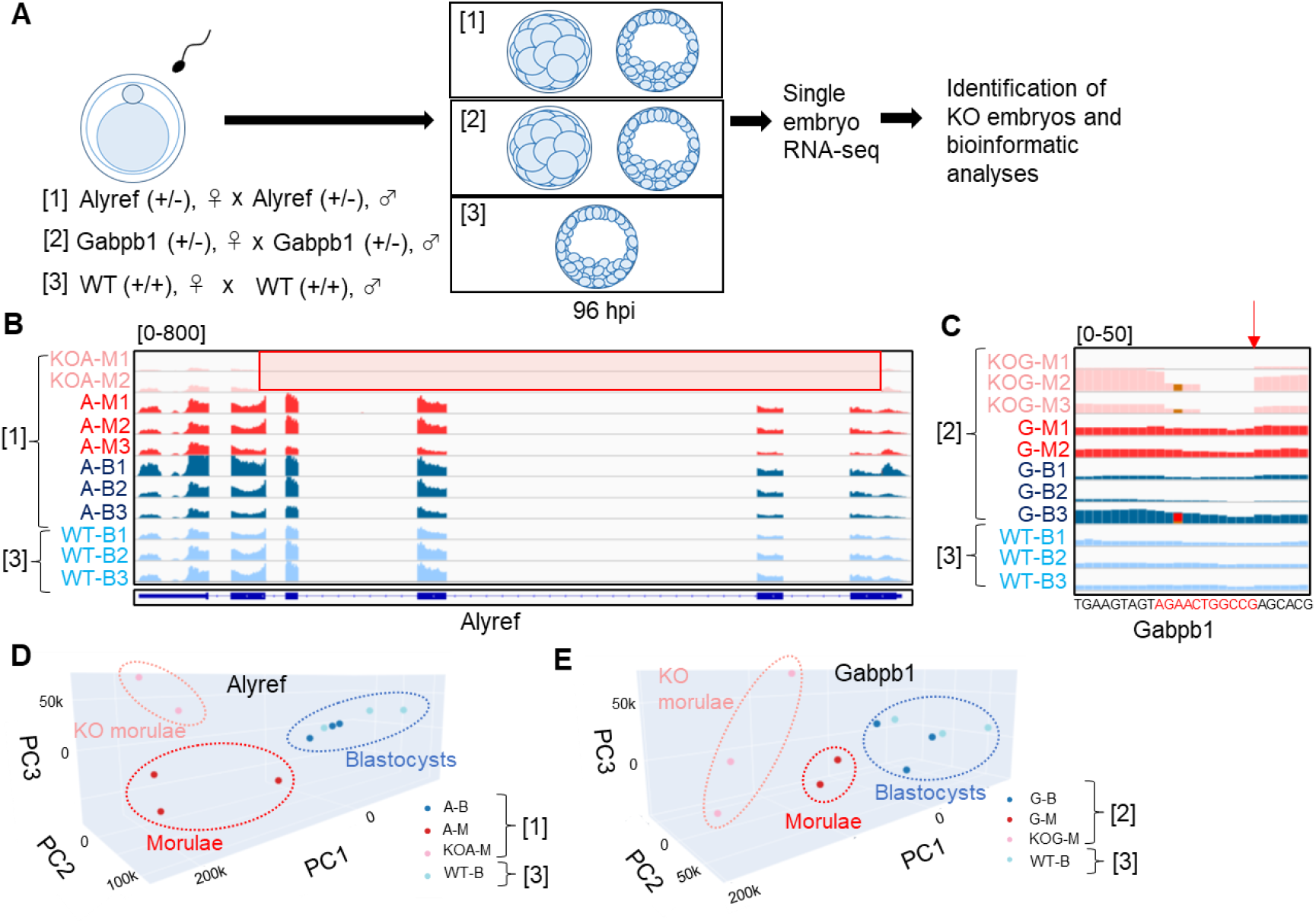
Abnormal transcriptomic patterns in *Alyref*^-/-^ or *Gabpb1*^-/-^ embryos. (A) Experimental design to examine transcriptomes of *Alyref*^-/-^ or *Gabpb1*^-/-^ embryos. Three different combinations of IVF experiments were performed (1-3), and morula or blastocyst embryo was collected from each IVF experiment for RNA-seq analyses. (B and C) Track images of RNA-seq reads at *Alyref* and *Gabpb1* gene loci. Figure 5B shows sequence reads of morulae and blastocysts collected after fertilization between *Alyref* heterozygous mice or control WT mice as shown in Figure 5A ([1] and [3]), while Figure 5C represents those collected after fertilization between *Gabpb1* heterozygous mice or control WT mice as shown in Figure 5A ([2] and [3]). *Alyref*^-/-^ embryos (KOA-M1, 2) were recognized by the lack of sequence reads at the deleted region (red highlighted box). Non-knockout morula and blastocyst embryos obtained from fertilization between *Alyref* heterozygous mice are shown as A-M1-3 and A-B1-3, respectively, while control WT blastocyst embryos are named as WT-B1-3. Similarly, *Gabpb1*^-/-^ embryos (KOG-M1-3) were identified due to the lack of sequence reads from the deleted locus (red arrow). Non-knockout morula and blastocyst embryos obtained from fertilization between *Gabpb1* heterozygous mice are shown as G-M1,2 and G-B1-3, respectively (D and E) Principal-component analysis (PCA) of the global gene expression profile among different samples after fertilization between *Alyref* heterozygous mice (Figure 5D) and that between *Gabpb1* heterozygous mice (Figure 5E). WT blastocyst embryos are used as a control (WT-B). PC1–3, principal components 1–3.

We next examined molecular pathways that were disturbed by the depletion of *Alyref*. DEGs were identified between knockout and wild type (and/or heterozygous) morulae (KOA- M1-2 vs A-M1-3) and 705 genes were misregulated (p < 0.05, 2-fold changes). Ingenuity pathway analysis (IPA) using this DEG list showed that genes related to mammalian pluripotency were misregulated (Figure 6A). We then searched for upstream regulators responsible for the abnormal gene expression in Alyref^-/-^ morulae using IPA. Interestingly, Pou5f1, a master regulator for mammalian pluripotency, was identified (Figure 6B). We examined Pou5f1-positive cells in *Alyref* KO embryos. Expansion of Pou5f1-positive cells and the proper formation of inner cell mass (ICM) were not observed in Alyref-negative embryos (Figure 6C). In addition, Pou5f1 signals were significantly reduced in *Alyref* knockdown embryos (Figure S6A, p < 0.01). These results suggest that Alyref might work as an upstream regulator or co-factor of Pou5f1 in early embryos. We tested this idea by overexpressing *EGFP- Alyref* in one blastomere of 2-cell embryos and asked if the overexpression of *EGFP-Alyref* enhanced the cell lineage contribution to ICM. We then counted the ratio of Pou5f1-expressing ICM cells in EGFP-Alyref-positive cells at the blastocyst stage. In *EGFP-Alyref* mRNA-injected embryos, 50% of EGFP-Alyref-positive cells contributed to ICM, while 39% of control histone H2B-EGFP-injected cells were found in the ICM (Figure 6D, p < 0.05). These results suggest that *Alyref* enhances the formation of ICM and is required for the developmental progression to blastocyst embryos.

**Figure 6.**
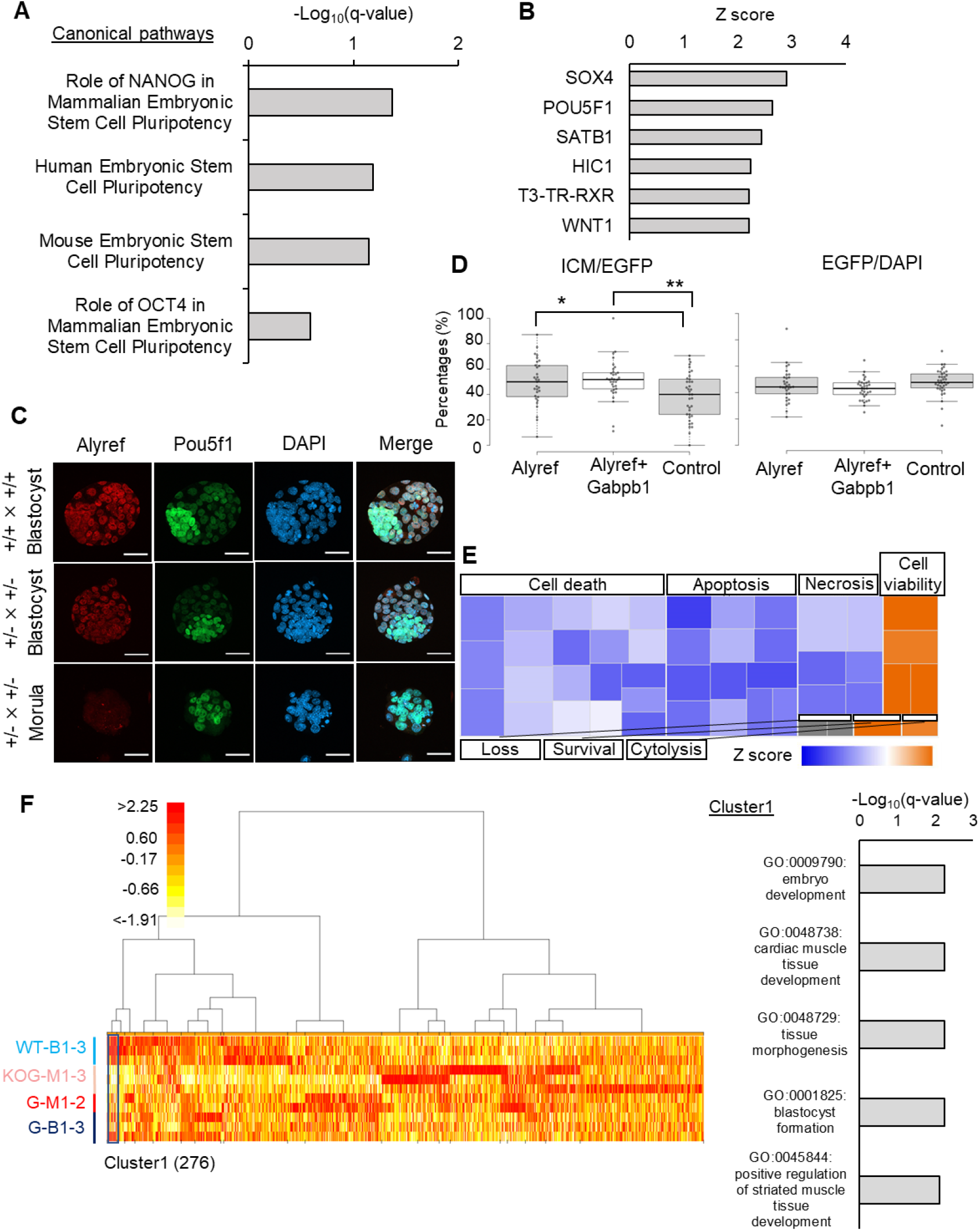
Gene expression programs related to the formation of blastocysts are impaired in *Alyref*^-/-^ and *Gabpb1*^-/-^ embryos. (A) Canonical pathways predicted by IPA using DEGs between knockout and wild type (and/or heterozygous) morulae (KOA-M1-2 vs A-M1-3). Significant terms related to mammalian pluripotency are indicated. (B) Predicted upstream regulators responsible for misexpression in *Alyref*^-/-^ embryos were identified by IPA. Activation Z scores are shown. (C) Immunostaining of Alyref and Pou5f1 in IVF embryos derived from different combinations of crossing. Some of the arrested morula embryos were devoid of Alyref signals (*Alyref*^+/-^ x *Alyref*^+/-^). The numbers of embryos used for experiments are as follows; (*Alyref*^+/+^ x *Alyref*^+/+^): 72, and (*Alyref*^+/-^ x *Alyref*^+/-^): 95. Three independent experiments were performed. Scale bars, 50 μm. (D) Percentages of Pou5f1-positive cells among EGFP-Alyref-positive cells in blastocysts. *EGFP-Alyref, EGFP-Alyref* + *EGFP-Gabpb1*, or *histone H2B-EGFP* mRNAs were injected into one blastomere of a 2-cell stage embryo and the lineage contribution of the injected cells to ICM were examined. Statistical significance was calculated by Tukey HSD test (*p < 0.05, **p < 0.01). The numbers of embryos used for experiments are as follows; *Alyref*: 31, *Alyref* + *Gabpb1*: 33 and Control: 37. Three independent experiments were performed. (E) Genes related to the cellular functions were misregulated in *Gabpb1*^-/-^ embryos, as revealed by IPA. A heatmap of activation Z scores is shown. (F) A heatmap of gene expression levels after clustering based on expression patterns. Twenty clusters were generated and cluster 1 is indicated. Color key indicates log2FC(FoldChange). Gene ontology (GO) terms enriched in cluster 1 are also shown.

DEGs were identified between *Gabpb1* knockout and wild type morulae (KOG-M1-3 vs G-M1-2) and 1321 genes were misregulated (p < 0.05, 2-fold changes). IPA revealed that genes related to cell viability and apoptotic pathways showed abnormal expression (Figure 6E). Moreover, anti-oxidation and metabolic pathways were abnormally regulated in Gabpb1^-/-^ embryos (Figure S6B, highlighted in green). Candidate upstream regulators related to this misexpression were identified (Figure S6C). We then performed unsupervised clustering of genes based on their expression levels and 20 clusters were generated (Figure 6F). Genes that belong to cluster 1 showed very low expression in Gabpb1^-/-^ morulae, but were expressed in other samples with strongest expression in wild type blastocysts (Figure 6F). Cluster 1 was enriched with genes related to blastocyst formation, and thus knockout of *Gapbp1* leads to the incomplete activation of such developmentally important genes. In summary, the roles of *Alyref* and *Gabpb1* in mouse preimplantation development are different, but both genes are required for the establishment of proper gene expression programs for blastocyst formation.

### Preimplantation development of SCNT embryos is rescued by exogenous supplementation of *Alyref* and *Gabpb1*

We finally asked if the insufficient expression of *Alyref* and *Gabpb1* is a reason for the low developmental ability of SCNT embryos. *EGFP-Alyref* and *EGFP-Gabpb1* mRNA was injected to SCNT embryos reconstructed with cumulus cells and subsequence preimplantation development was examined (Figure 7A, without TSA). Since the developmental rescue is often dose-dependent, different doses of mRNA mixtures of *Alyref* and *Gabpb1* were injected (100, 400, 500, and 600 ng/μl). *Alyref* and *Gabpb1* mRNA injection enhanced preimplantation development of SCNT embryos (Figure 7B) and the most prominent effect was observed when 400 ng/μl of *Alyref* and *Gabpb1* mRNA (each 200 ng/μl) was injected (Figures 7B and 7C). As shown in Figure S2, *Alyref* and *Gabpb1* are activated by removing H3K9me3 with Kdm4d or lowing it with VC. We therefore hypothesized that the positive effect of VC on reprogramming in SCNT embryos can be substituted by the supplementation of *Alyref* and *Gabpb1* mRNA. VC’ s effect is most prominent when combined with TSA (Miyamoto et al., 2017) and therefore we injected 400 ng/μl of *Alyref* and *Gabpb1* mRNA to SCNT embryos treated with TSA (Figure 7A, with TSA). Strikingly, 87% of SCNT embryos developed to the blastocyst stage after *Alyref* and *Gabpb1* mRNA injection (Figure 7D). Taken together, improper activation of *Alyref* and *Gabpb1* hampers preimplantation development of SCNT embryos.

**Figure 7.**
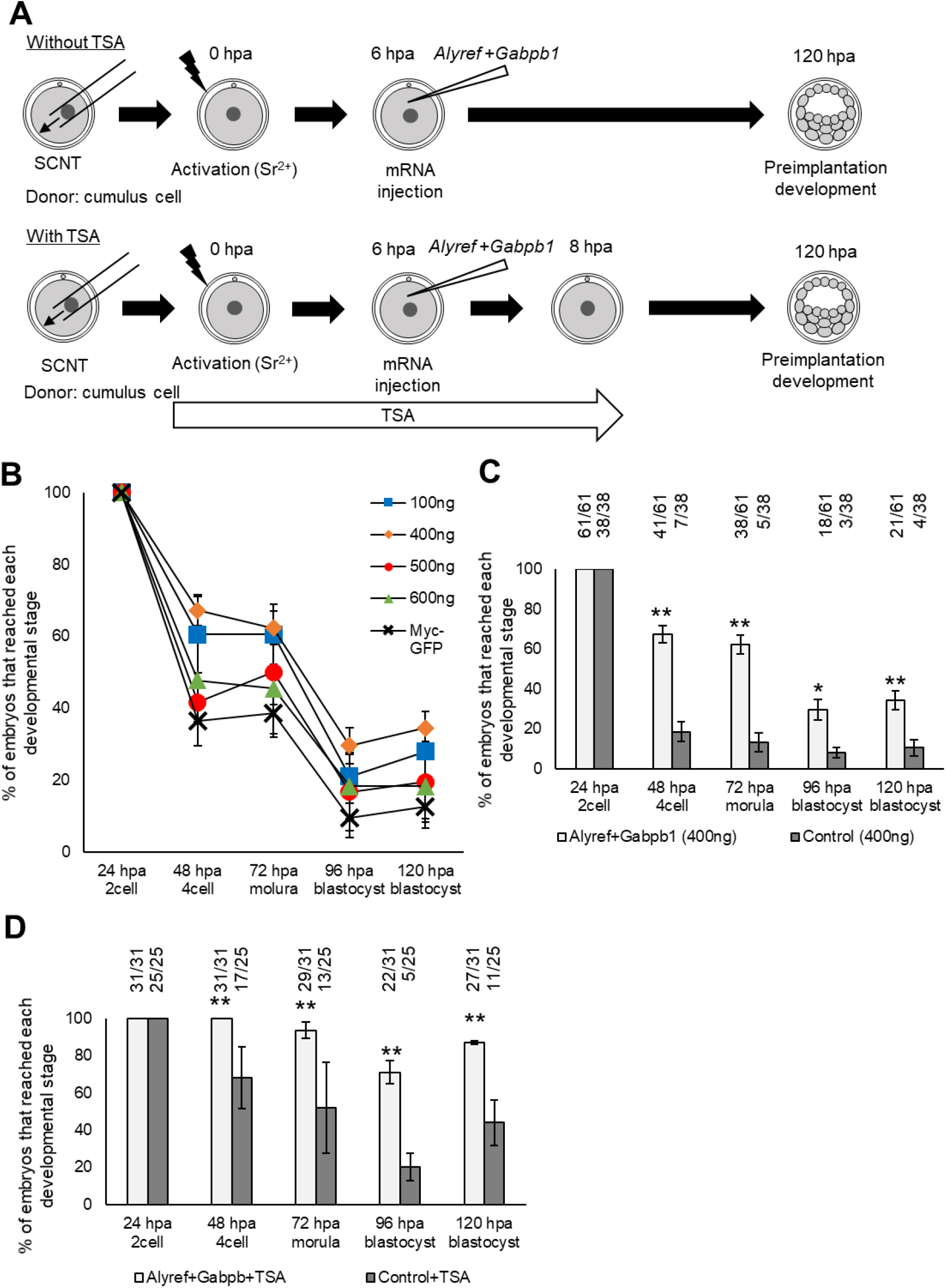
Developmental rescue of SCNT embryos by the supplementation of Alyref and Gabpb1. (A) Experimental design to examine the effect of *Alyref* and *Gabpb1* mRNA injection on the development of SCNT embryos with or without TSA treatment. hpa: hours post activation. (B) Preimplantation development of SCNT embryos injected with various different concentrations of *EGFP-Alyref* and *EGFP-Gabpb1* mRNA (100-600 ng/μl). *Myc-tag-GFP* mRNA (100∼600 ng/μl) was injected as a control. Error bars are SE. Three to Four independent experiments were performed. (C) Preimplantation development of SCNT embryos injected with 400 ng/μl of *EGFP-Alyref* and *EGFP-Gabpb1* mRNA or *Myc-tag-GFP* mRNA as a control. Error bars are SE. Statistical significance was calculated by chi-square test (*p < 0.05, **p < 0.01). The numbers of embryos used for experiments and that reached each developmental stage are indicated. Four independent experiments were performed. (D) Preimplantation development of TSA-treated SCNT embryos injected with 400 ng/μl of *EGFP-Alyref* and *EGFP-Gabpb1* mRNA or *Myc-tag-GFP* mRNA as a control. Error bars are SE. Statistical significance was calculated by chi-square test (**p < 0.01). The numbers of embryos used for experiments and that reached each developmental stage are indicated. Three independent experiments were performed.

## Discussion

Tremendous efforts have been made to improve the developmental potential of cloned embryos. Recent studies have revealed that the alteration of epigenetic states in SCNT embryos greatly enhances their development to term. Especially, the removal of the repressive H3K9me3 mark is key for development. However, it is still unclear how the reduced H3K9me3 helps development of SCNT embryos. One assumption is that genes necessary for embryonic development are fully activated upon the removal of H3K9me3 because the overexpression of H3K9me3 demethylase Kdm4d upregulated expression of many zygotically activated genes (Matoba et al., 2014). In line with this concept, our study demonstrates that expression of H3K9me3-regulated *Alyref* and *Gapbp1* is normally inhibited in SCNT embryos, and the exogenous supplementation of such mRNA can rescue the development of SCNT embryos. Furthermore, we have shown that these genes are necessary for preimplantation development of fertilized embryos. Thus, our paper provides mechanistic insight into somatic cell nuclear reprogramming in SCNT embryos and further represents a valid approach to identifying developmentally required genes.

Our siRNA screening identified two developmentally important genes *Alyref* and *Gabpb1*. Embryonic lethal phenotype till E12.5 has been reported after knocking out *Gapbp1*(Xue et al., 2008), but its role for preimplantation development has not been investigated. Our study shows that knockout of *Gabpb1* induces the abnormal gene expression pattern, especially to genes related to antioxidation and cellular viability. GABPB1 forms a tetrameric complex and activates the transcription of various genes, including the antioxidant genes (Usmanova et al., 2011), suggesting the defective expression of key metabolic genes may lead to the arrested phenotype around the morula stage. Strikingly, all embryos were arrested at the morula stage after knocking out *Alyref*. Alyref has been shown as a reader protein for 5- Methylcytosine (5mC) RNA and involved in the processing and export of such RNA (Yang et al., 2017). 5mC modification facilitates the mRNA stability during maternal to zygotic transition in zebrafish (Yang et al., 2019). In our study, the inhibition of *Alyref* caused insufficient expression of Pou5f1 protein. It is therefore plausible that Alyref might be involved in the dynamics of *Pou5f1* mRNA. It is also noteworthy that the overexpression of Alyref in one of the blastomeres at the 2-cell stage enhanced the cell lineage contribution to ICM (Figure 6D). Future studies need to clarify whether Alyref directly enhanced Pou5f1 expression for ICM specification or Alyref worked cooperatively with Pou5f1 for blastocyst formation. Interestingly, defective activation of Pou5f1 in SCNT embryos is often observed (Boiani et al., 2002). Insufficient activation of Alyref in SCNT embryos might be related to this phenotypic defect of SCNT embryos.

Repressive histone marks serve as a barrier for somatic cell nuclear reprogramming. *Alyref* and *Gabpb1* were identified as genes upregulated upon the removal of H3K9me3 and those by VC treatment that also lowers H3K9me3 levels (Figure S2); otherwise repressed in SCNT embryos (Figure 2). At least two different routes can be considered for the mechanism of silencing *Alyref* and *Gabpb1* by H3K9me3 in SCNT embryos. Firstly, expression of *Alyref* and *Gabpb1* is directly inhibited by H3K9me3, in which case the forced removal of H3K9me3 in donor cells should allow activation of *Alyref* and *Gabpb1*. Secondly, transcriptional activators of *Alyref* and *Gabpb1* are suppressed by H3K9me3, eventually resulting in the repression of *Alyref* and *Gabpb1*. In the later case, it might be difficult to alter chromatin states of donor cells for expression of *Alyref* and *Gabpb1* in SCNT embryos. Instead, supplementation of *Alyref* and *Gabpb1* mRNA to SCNT embryos by microinjection would rescue their expression in both cases and we have shown that the preimplantation development of SCNT embryos can be enhanced by injecting *Alyref* and *Gabpb1* mRNA (Figure 7). These results suggest that incomplete activation of *Alyref* and *Gabpb1* is at least partially responsible for the low developmental potential of SCNT embryos. However, considering that the remarkable rescue of development to blastocysts is only observed in combination with TSA (Figure 7D), insufficient expression of *Alyref* and *Gabpb1* cannot fully explain the H3K9me3-caused low developmental ability of SCNT embryos. Further investigations are needed for the comprehensive understanding of the H3K9me3-mediated barrier for nuclear reprogramming to the totipotent state.

Based on these observations, we propose the nature of the low developmental ability of SCNT. Firstly, the condensed chromatin state of somatic nuclei prevents the access of maternal reprogramming factors. Global relaxation of chromatin by increasing histone acetylation, such as by TSA, overcomes this barrier. Secondly, reprogramming-resistant H3K9me3 marks in somatic chromatin prevents activation of developmental genes. Our study demonstrates that *Alyref* and *Gabpb1* are one of the key genes that are normally inhibited at the time of ZGA in SCNT embryos by the effect of reprogramming-resistant H3K9me3. Considering the fact that overexpression of H3K9me3 demethylases or TSA+VC treatment can more efficiently enhance development of SCNT embryos than *Alyref* and *Gabpb1* overexpression, other unidentified developmental genes seem to be also repressed by H3K9me3. Thirdly, the lack of H3K27me3- mediated imprinting in somatic chromatin results in misregulation of miRNA clusters, which are closely related to the proper placenta formation (Inoue et al., 2020; Wang et al., 2020). Taken together, it is intriguing to know that the impaired development of SCNT embryos can be rescued by several key factors at different developmental stages, indicative of progressive reprogramming in SCNT embryos. Our study sheds light on the previously unexplored genes that are important for somatic nuclei to obtain totipotency after nuclear transfer. Furthermore, we show the proof of concept that impaired preimplantation development of SCNT embryos is caused by the incomplete activation of developmentally necessary genes at ZGA.

## ACKNOWLEDGEMENTS

We thank Mr Kajikuri, Ms Matsuzawa, and Ms. Kusakabe for their contributions to initiating the project. We thank K.K. DNAFORM (Yokohama, Japan) for RNA-seq analyses. We thank Ms N Backes Kamimura for proofreading. This research was supported by JSPS KAKENHI Grant Numbers JP19H05271, JP19H05751, JP20K21376 to K.Miyamoto. by The Naito Foundation to K.Miyamoto, by Takeda Science Foundation to K.Miyamoto, by a Kindai University Research Grant (19-II-1) to K.Miyamoto.

## AUTOR CONTRIBUTIONS

K.Miyamoto conceived the project. K.Miyamoto designed experiments. S.I., M.H., M.K., M.M., Y.I., M.N., M.A., and K.Miyamoto performed experiments. K.Matsumoto and M.I. provided essential materials. S.I., M.N., and K.Miyamoto analyzed imaging data. K.Miyamoto performed bioinformatics analyses. M.I. and K.Miyamoto supervised research. S.I. and K.Miyamoto wrote the manuscript.

## Methods

### RESOURCE AVAILABILITY

#### Lead Contact

Further information and requests for resources and reagents should be directed to and will be fulfilled by the Lead Contact, Kei Miyamoto.

## Materials Availability

All unique/stable reagents generated in this study are available from the Lead Contact with a completed Materials Transfer Agreement.

## Data and Code Availability

The dataset generated during this study are available in the Gene Expression Omnibus (GEO) public repository under accession GEO: GSE199874. All the other data are available from the Lead Contact upon reasonable request.

## EXPERIMENTAL MODEL AND SUBJECT DETAILS

### Animals

Mice (C57BL/6, DBA/2 strains, and B6D2F1 (C57BL/6J×DBA/2N)) at 8-10 weeks of age were purchased from CLEA Japan (Tokyo, Japan) or Japan SLC (Shizuoka, Japan) and maintained in light-controlled, air-conditioned rooms. B6D2F1 mice were used for *in vitro* fertilization. This study was carried out in strict accordance with the recommendations in the Guidelines of Kindai University for the Care and Use of Laboratory Animals. Experimental protocols were approved by the Committee on the Ethics of Animal experiments of Kindai University (Permit Number: KABT-31-003). All mice were sacrificed by cervical dislocation and all efforts were made to minimize suffering and to reduce the number of animals used in the present study.

KO mice were generated on the B6D2F1 background with CRISPR/Cas9-mediated genome editing as reported before (Mashiko et al., 2013; Oji et al., 2016). Genotyping of KO mice was performed as described in Figures 3 and 4. PCR products were directly used for Sanger sequencing. Sequences of gRNA and primers are listed in Table S1. Animal experiments for generating KO mice were approved by the Animal Care and Use Committee of the Research Institute for Microbial Diseases, Osaka University (Permit Number: H30-01-1 and R03-01-0).

## METHOD DETAILS

### In Vitro Fertilization (IVF) and Embryo Culture

Collection of spermatozoa, oocytes, and zygotes were performed as described in previous studies (Okuno et al., 2020). Briefly, spermatozoa were collected from the cauda epididymis of B6D2F1 fertile male mice (>8 weeks of age). The sperm suspension was incubated in human tubal fluid (HTF) medium for 1.5 hours to allow for capacitation at 37°C under 5% CO_2_ in air. Oocytes were collected from the excised oviducts of B6D2F1female mice (>8 weeks of age) that were superovulated with pregnant mare serum gonadotropin (PMSG; Serotropin, ASKA Pharmaceutical Co., Tokyo, Japan) and 48 hours later, human chorionic gonadotropin (hCG; ASKA Pharmaceutical Co.). Cumulus-oocyte complexes were recovered into pre-equilibrated HTF medium. The sperm suspension was added to the oocyte cultures and morphologically normal zygotes were collected 2 hours post insemination (hpi). The zygotes were cultured in potassium simplex optimized medium KSOMaa (ARK Resource, Kumamoto, Japan) at 37°C under 5% CO_2_ in air.

### Somatic cell nuclear transfer (SCNT)

SCNT was carried out as described previously (Miyamoto et al., 2017). Briefly, enucleation of denuded MII oocytes was performed in drops of HCZB containing 5 µg/ml cytochalasin B (Sigma-Aldrich). After enucleation, a donor cell in HCZB with 6% dBSA was placed in the perivitelline space of an enucleated oocyte together with HVJ-E (GenomeONE-CF, Ishihara Sangyo, Osaka, Japan) by tightly attaching the donor cell to the enucleated oocyte, the oocyte was then cultured in KSOMaa for 1 h at 37°C in air containing 5% CO_2_, during which time it fused with the donor cell. The reconstructed oocytes were activated by the incubation for 6 h in 5 mM SrCl_2_ and 2 mM EGTA-containing KSOMaa supplemented with 5 µg/ml cytochalasin B, referred to as activation medium (Kishigami and Wakayama, 2007), at 37°C in air containing 5% CO_2_. For the treatment with trichostatin A (TSA, Sigma-Aldrich, cat. code: T8552), NT embryos were treated with 50 nM TSA for 8 h from the commencement of activation. The activated NT embryos were cultured at 37°C in air containing 5% CO_2_ in KSOMaa.

### Plasmids

Constructs were produced either using a gateway system (Thermo Fisher Scientific) or in-fusion cloning (Clontech). Myc-tag-GFP was subcloned into the pCS2 vector (Miyamoto et al., 2011) for mRNA production as described previously (Baarlink et al., 2017). pCS2-myc-GFP and pcDNA3.1-Histone H2B-EGFP were reported previously (Baarlink et al., 2017; Hatano et al., 2022). To obtain the *Gabpb1* mRNA producing vectors for injection experiments, we first generated the pENTR/D-TOPO-Gabpb1 entry vector. After clonase-triggered recombination reaction, pCS2-EGFP-Gabpb1 was generated. *Gabpb1* insert was amplified from cDNA of mouse blastocyst embryos by using oligos GW_Gabpb1_N_F and GW_Gabpb1_N_R (Table S1). pCS2-EGFP-Alyref was produced by in-fusion cloning. For this subcloning, *Alyref* insert was amplified from the pCMV6-Entry-Alyref-Myc-DDK-Tagged plasmid (OliGene Technologies, Rockville, USA, MR220236) using IF-Alyref-N_Fw and IF-Alyref-N_Rv3 primers. In-fusion reaction was performed after amplification of the linearized pCS2 vector, which was amplified by PCR using IF-Alyref-vectorF and IF-Alyref-vectorR primers. All primers used for cloning are listed in Table S1.

### mRNA Production

mRNAs were prepared from pCS2 vectors using mMESSAGE mMACHINE SP6 Transcription Kit (Thermo Fisher Scientific, AM1340 or AM1344). Briefly, to produce linearized vectors, approximately 5 μg plasmids were digested overnight, with appropriate restriction enzymes. In the case of pCS2 vectors, mRNAs produced from the SP6 promoter were subjected to the addition of polyA tails (Thermo Fisher Scientific, AM1350). Produced mRNAs were purified using Rneasy Mini Kit (QIAGEN, 74106).

### mRNA Injection

After activation, NT embryos were collected at 6∼7 hours post activation (hpa) for mRNA injection. NT embryos were then washed with KSOMaa and kept in drops of CZB-HEPES medium for injection. mRNAs were injected using a piezo manipulator (Prime Tech, Tsukuba, Japan). The final concentrations of injected mRNA are as follows; 50-300 ng/μl *EGFP-Alyref* mixed with 50-300 ng/μl *EGFP-Gabpb1*, 100-600 ng *Myc-tag-GFP* for Figure 7B, 200 ng/μl *EGFP-Alyref* mixed with 200 ng/μl *EGFP-Gabpb1*, 400 ng *Myc-tag-GFP* for Figures 7C and D. After injection, embryos were cultured in KSOMaa medium at 37°C in air containing 5% CO2. For the TSA treatment, 50 nM TSA was supplemented in KSOMaa medium and NT embryos were cultured in the TSA-containing medium. After injecting mRNA, zygotes were immediately transferred to the inhibitor-containing medium.

For Figure 6D, IVF embryos were collected at 24∼27 hpi for mRNA injection. Injection to 2-cell embryos was performed in drops of CZB-HEPES medium. mRNAs were injected into one blastomere of a two cell embryo using a piezo manipulator. The final concentrations of injected mRNA are as follows; 50 ng/μl *EGFP-Alyref*, 50 ng/μl *EGFP-Alyref* mixed with 50 ng/μl *EGFP-Gabpb1*, 50 ng *Histone H2B-EGFP* for Figure 6D. After injection, embryos were cultured in KSOMaa medium at 37°C in air containing 5% CO_2_.

### Immunofluorescence Staining

Embryos were fixed in 4% PFA/PBS at room temperature for 15 min, and were washed by 3 mg/ml PVP/PBS for 3 times. Zona pellucida was removed in Tyrode’ s solution (SIGMA-Aldrich, T1788) and the zona-free embryos were washed by 3 mg/ml PVP/PBS for 3 times. Samples were next treated with 0.25% Triton X-100 in 3 mg/ml PVP/PBS at room temperature for 30 min. Blocking was performed in 3% BSA/PBS with 0.01% Tween 20 for 1 hour at room temperature, then embryos were incubated with primary antibodies diluted in 3% BSA/PBS with 0.01% Tween 20 (1:50; Anti-ALY Antibody [sc-32311, Santa Cruz Biotechnology], 1:200; GABPB1 antibody [GTX103464, Gene Tex], 1:200; Oct4 Antibody [09-0023, REPROCELL], 1:100; Oct4 Antibody [sc5279, Santa Cruz Biotechnology]) at 4°C overnight. Following three times washes by 1% BSA/PBS with 0.01% Tween 20, samples were further incubated in the dark with Alexa Fluor 488-labeled goat anti-mouse IgG antibody (1:2,000; A11001, Thermo Fisher Scientific), Alexa Fluor 488-labeled goat anti-rabbit IgG antibody (1:2,000; A11008, Thermo Fisher Scientific), Alexa Fluor 594-labeled donkey anti-mouse IgG antibody (1:2,000; A21203, Thermo Fisher Scientific), or Alexa Fluor 594-labeled donkey anti-rabbit IgG antibody (1:2,000; A21207, Thermo Fisher Scientific) at room temperature for 1 hour. The samples were washed with 1% BSA/PBS with 0.01% Tween 20 three times and then mounted on slides using VECTASHIELD Mounting Medium containing DAPI. The fluorescence signals were observed using a LSM800 microscope, equipped with a laser module (405/488/561/640 nm) and GaAsP detector, using the same contrast, brightness, and exposure settings within the same experiments. Z-slice thickness was determined by using the optimal interval function in the ZEN software.

### Image Analysis

Images were analyzed using the ZEN software. For quantification of Alyref signals in SCNT embryos (Figure 2C), all focal planes were merged by maximum intensity projection. Alyref staining did not show non-specific cytoplasmic signals and it was localized in nuclei. Therefore, the maximum intensity projection using all focal planes was used for quantifying Alyref signals. After maximum intensity projection, intensities of Alyref in nuclei was quantified. Furthermore, background signals were subtracted from the nuclear signals and the average signal intensities of merged focal planes were calculated by using the “Measure” function in the ZEN software. In order to quantify nuclear signal intensities of Gabpb1 (Figure 2D), all focal planes that cover the nuclear region were used for quantifying nuclear signal intensities. In each focal plane, background signals were subtracted from the nuclear signals and the average of all focal planes was calculated by using the “Measure” function in the ZEN software.

For Figure 6D, the injected embryos were cultured till 88 hpi and blastocyst embryos were collected. The collected blastocysts were fixed and stained with Pou5f1 and DAPI. Then, EGFP-positive cells and Pou5f1-positive ICM cells, together with the total number of cells with DAPI, were counted by the confocal microscopy.

### RNA-sequencing (RNA-seq) Analysis

Timings for sampling are shown in each corresponding figure. Single NT embryo at the morula or blastocyst stage was treated with acid Tyrode, followed by three times washes with 0.1% BSA/PBS, and were moved into 1×Reaction buffer from SMART-seq v4 Ultra Low Input RNA Kit (Z4888N, Takara). SMART-seq library preparation was performed using SMART-seq v4 Ultra Low Input RNA Kit and Nextera DNA Sample Preparation Kit (FC-131-1024, illumine, San Diego, CA) according to the vendor’ s instruction. We followed the previously published protocol for RNA-seq of mouse embryos (Tomikawa et al., 2021).

Briefly, paired-end sequencing (50 bp + 25 bp) was obtained by the NextSeq platform (Illumina). Raw reads were first subjected to filtering to remove low quality reads using Trimmomatic (Bolger et al., 2014). Reads of less than 20 bases and unpaired reads were also removed. Furthermore, adaptor, polyA, polyT and polyG sequences were removed using Trim Galore! (https://www.bioinformatics.babraham.ac.uk/projects/trim_galore/). The sequencing reads were then mapped to the mouse genome (mm10) using STAR (Dobin et al., 2013). Reads on annotated genes were counted using featureCounts (Liao et al., 2014). RNA-seq reads were visualized using Integrative Genome Viewer (Robinson et al., 2011). FPKM values were calculated from mapped reads by normalizing to total counts and transcript. Differentially expressed genes (DEGs, p < 0.05) were then identified using DESeq2 (Love et al., 2014). An unsupervised hierarchical clustering of the read count values was performed using hclust in TCC (unweighted pair group method with arithmetic mean: UPGMA). Gene ontology terms showing over-representation of genes that were up or downregulated were detected using DAVID tools (p < 0.05). Principal-component analysis (PCA) of the global gene expression profile was performed using scikit-learn and the plots were depicted using plotly. Each gene list was further subjected to an Ingenuity Pathway Analysis (IPA; QIAGEN, Redwood City, CA). Using IPA, enriched canonical pathways, upstream transcriptional regulators, and diseases and biological functions were investigated.

For Figures S2B and S2C, FPKM values were obtained from GSE59073 (Matoba et al., 2014) and GSE95053 (Miyamoto et al., 2017).

### RT-qPCR

For examining the effect of siRNA knockdown, total RNA was extracted from a pool of three 4- cell embryos using PicoPure RNA Isolation Kit (Life Technologies; KIT0204) according to the manufacturer’ s protocol. cDNA was synthesized from the extracted RNA using Superscript III RT First-Strand Synthesis System (Life Technologies; 18080051). cDNA samples were analyzed in 7300 Real Time PCR System (Applied Biosystems). Primer sequences are listed in Table S1.

### siRNA Knockdown

For investigating a function of candidate genes responsible for acquiring totipotency, specific siRNA was injected to IVF embryos at 6 hpi, and the injected embryos were further cultured in KSOMaa medium to observe preimplantation development. Pre-designed negative control siRNA (RNAi Inc., Tokyo, Japan) was used as a control. Sequences for siRNA and qPCR primers are listed in Table S1.

## QUANTIFICATION AND STATISTICAL ANALYSIS

All of the statistical methods for each experiment can be found in the figure legends as well as in the Method Details section. For Figures 1C, 2C, 2D, 4D, S1A, S3B, and S6A, statistical significance was calculated by two-sided T-test. For Figures 1D, 3C, 4E, 7C, 7D, and S1B, statistical significance was calculated by chi-square test. For Figure 6D, Tukey HSD test was used.

