## Supplementary information for "Incomplete activation of developmentally required genes *Alyref1* and *Gabpb1* leads to preimplantation arrest in cloned mouse embryos"

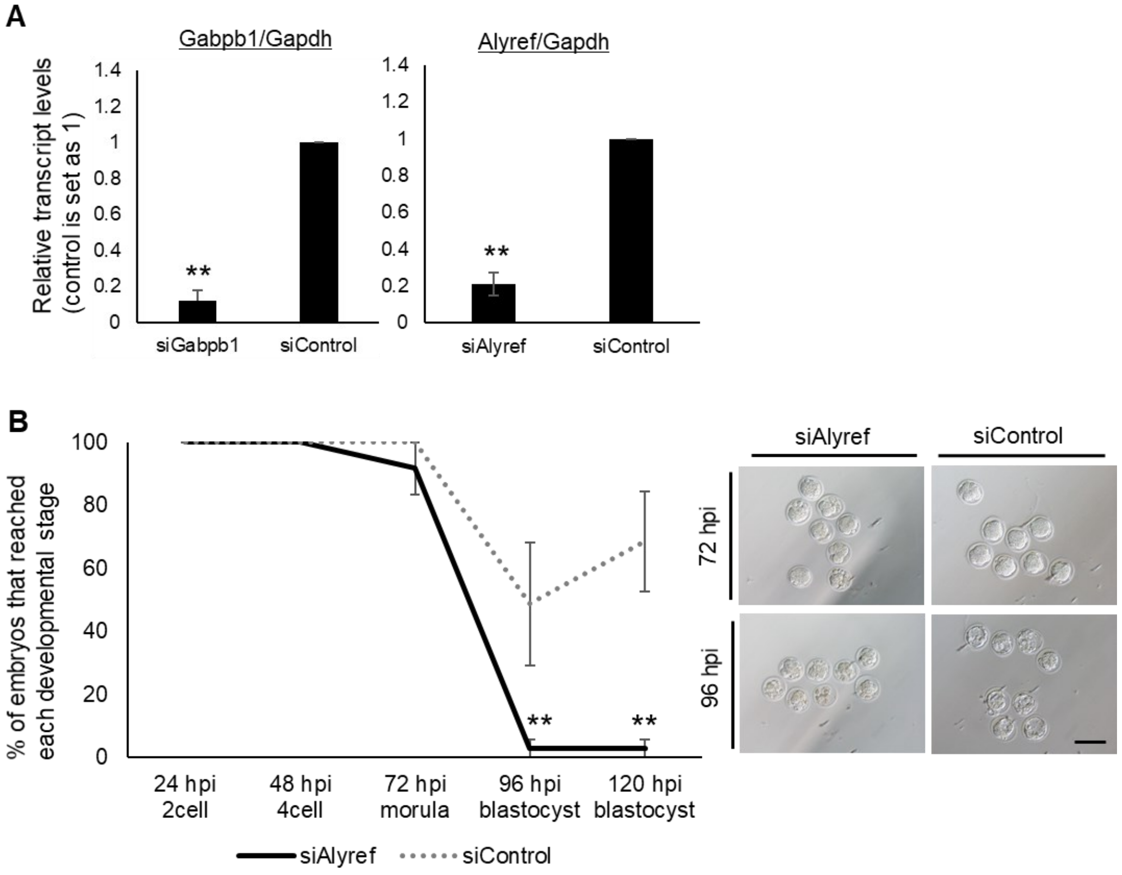


**Figure S1. Knockdown of *Alyref* and *Gabpb1* in mouse preimplantation embryos and development of such embryos, related to Figure 1**

(A) RT-qPCR analysis of *Alyref* siRNA-injected and *Gabpb1* siRNA-injected embryos at the 4-cell stage. Gene expression was normalized by *Gapdh* expression. Statistical significance was calculated by two-sided T-test. Three (*Gabpb1*) and four (*Alyref*) independent experiments were performed. Error bars are SE. **$p < 0.01$.

(B) Preimplantation development of embryos injected with siRNA against *Alyref* and control siRNA. Representative images of the injected embryos are shown in the right panel. Statistical significance was calculated by chi-square test. The numbers of embryos used for experiments are as follows: siAlyref: 32, and siControl: 25. Three independent experiments were performed. Error bars are SE. Scale bar, 100 µm. **$p < 0.01$.

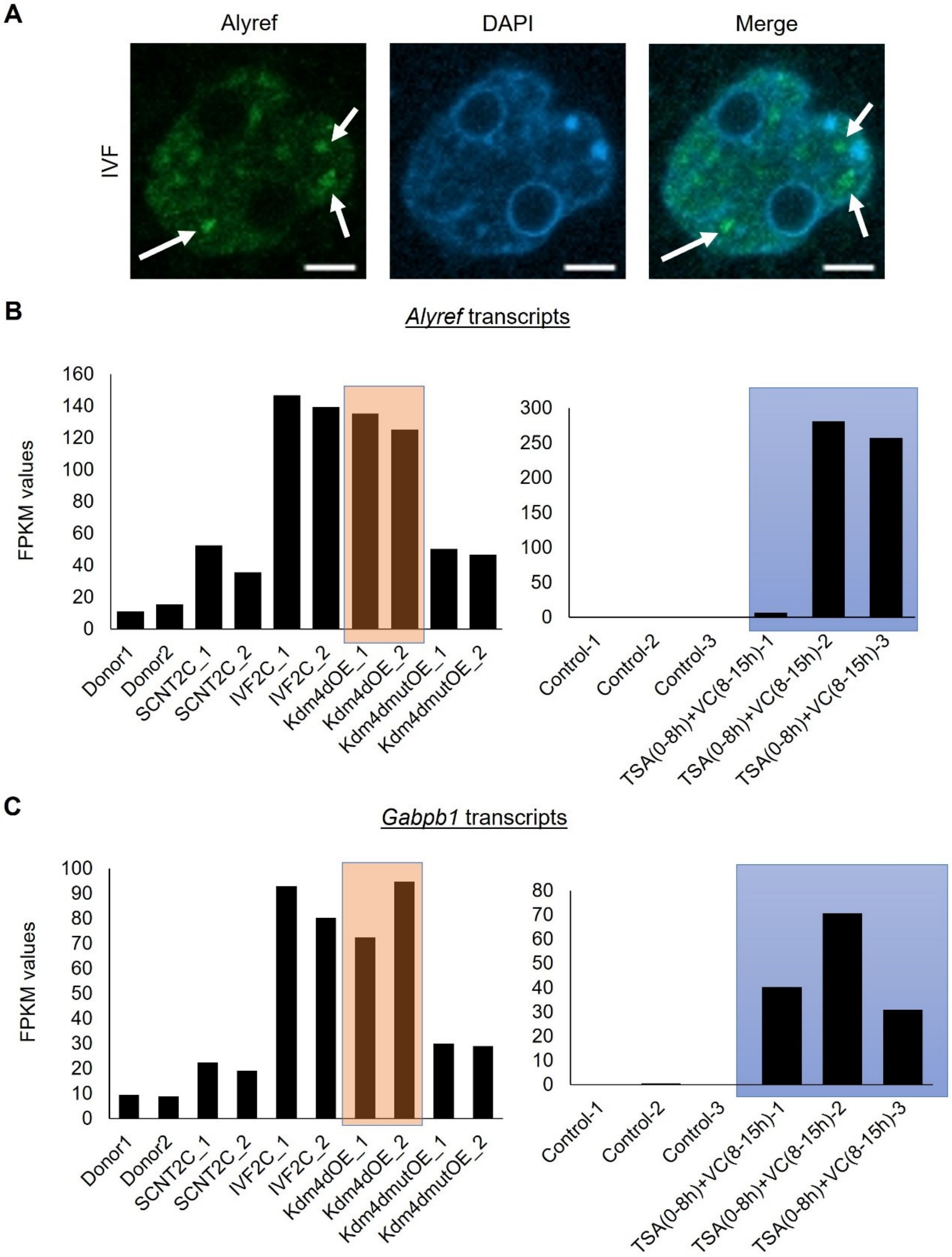


**Figure S2. Expression of Alyref and Gabpb1 in mouse embryos, related to Figure 2**

(A) Immunostaining of Alyref in fertilized embryos at the 2-cell stage and Alyref localizes as dots in nuclei (arrows). DNA was stained with DAPI. Merge represents the merged photo. n =

26. Three independent experiments were performed. Scale bars, 5 µm.

(B and C) Fragments per kilobase of exon per million mapped (FPKM) values of Alyref and Gabpb1 in donor cells, control IVF embryos and SCNT embryos treated with various conditions (Kdm4dOE: Kdm4d overexpression, Kdm4dmutOE: Kdm4d mutant overexpression, TSA(0-8h)+VC(8-15h): TSA+VC treatment) (Matoba et al., 2014; Miyamoto et al., 2017). Embryos are collected at the 2-cell stage and subjected to RNA-seq analyses. Controls in the right graphs represent SCNT embryos without TSA+VC treatment. Kdm4d overexpression in SCNT embryos rescues expression of Alyref and Gabpb1 (red box) to the level equivalent with IVF embryos, and treatment with TSA+VC upregulates expression of Alyref and Gabpb1in SCNT embryos (blue box).

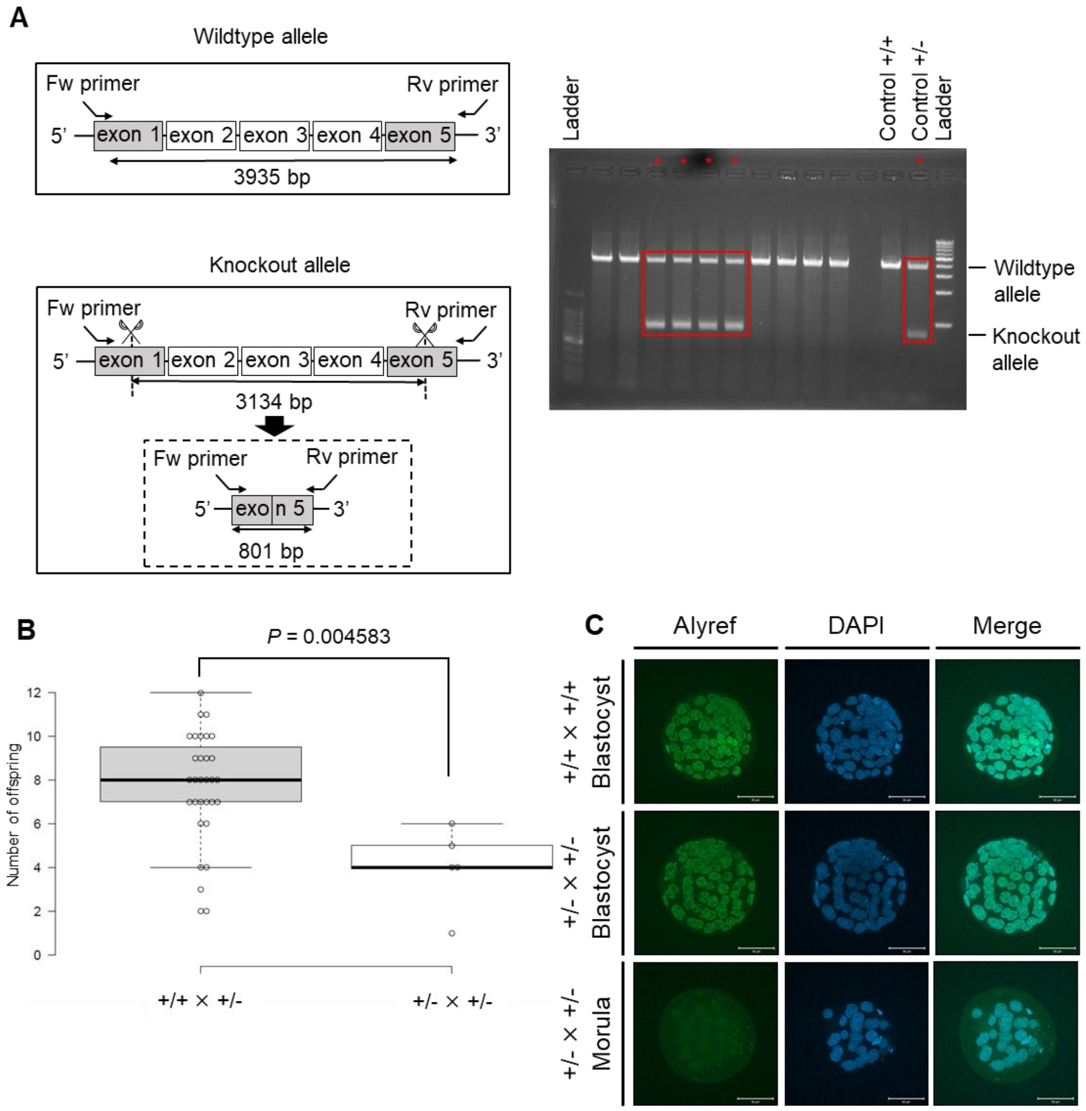


**Figure S3. Knockout of *Alyref* gene leads to the embryonic lethal phenotype, related to Figure 3**

(A) A scheme for PCR-mediated genotyping of *Alyref* gene. Primers were designed to span the knockout region (3134 bp). Positions of primers and the product sizes for wild type and knockout alleles are indicated (3935 bp and 801 bp, respectively). Genotyping results of Alyref$^{+/-}$ mice are highlighted by red boxes in the gel image.

(B) A box plot indicates the number of offspring after each mating. Statistical significance was calculated by two-sided T test.

(C) Immunostaining of Alyref in IVF embryos derived from different combinations of crossing.

Some of the arrested morula embryos were devoid of Alyref signals (+/- x +/-). The numbers of embryos used for experiments are as follows: (+/+ x +/+): 144, and (+/- x +/-): 74. Four independent experiments were performed. Scale bars, 50 μm.

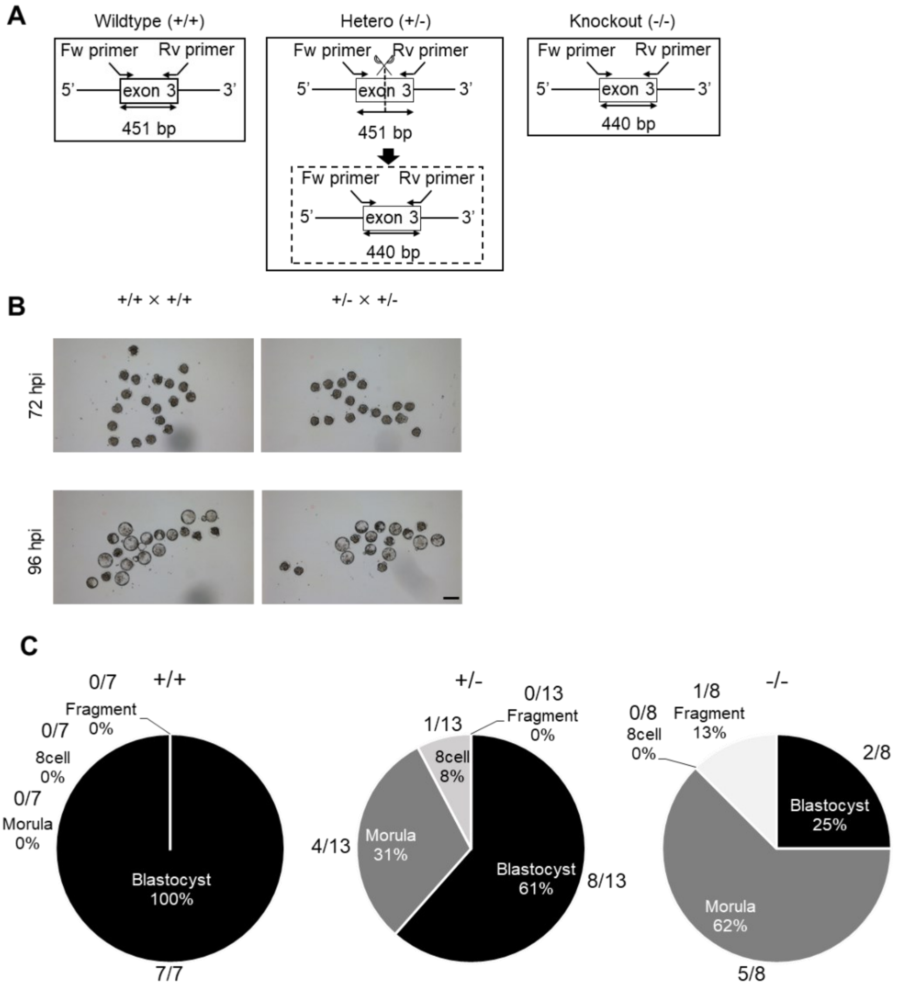


**Figure S4. Knockout of *Gabpb1* gene leads to the embryonic lethal phenotype, related to Figure 4**

(A) PCR primers that detect both wild type and knockout *Gabpb1* alleles for genotyping. Primers were designed to span the deleted region. Positions of primers and each genotype product size are indicated.

(B) Representative images of IVF embryos. Control Gabpb1$^{+/+}$ x Gabpb1$^{+/+}$ embryos are shown in left panels, while Gabpb1$^{+/-}$ x Gabpb1$^{+/-}$ embryos are shown in right. Scale bar, 100 µm.

(C) Pie charts represent the breakdown of developmental stages in IVF-derived Gabpb1$^{+/+}$ (left), Gabpb1$^{+/-}$ (middle), and Gabpb1$^{-/-}$ embryos (right) at 96 hpi. The numbers of embryos are also shown. Fragment represents fragmented embryos.

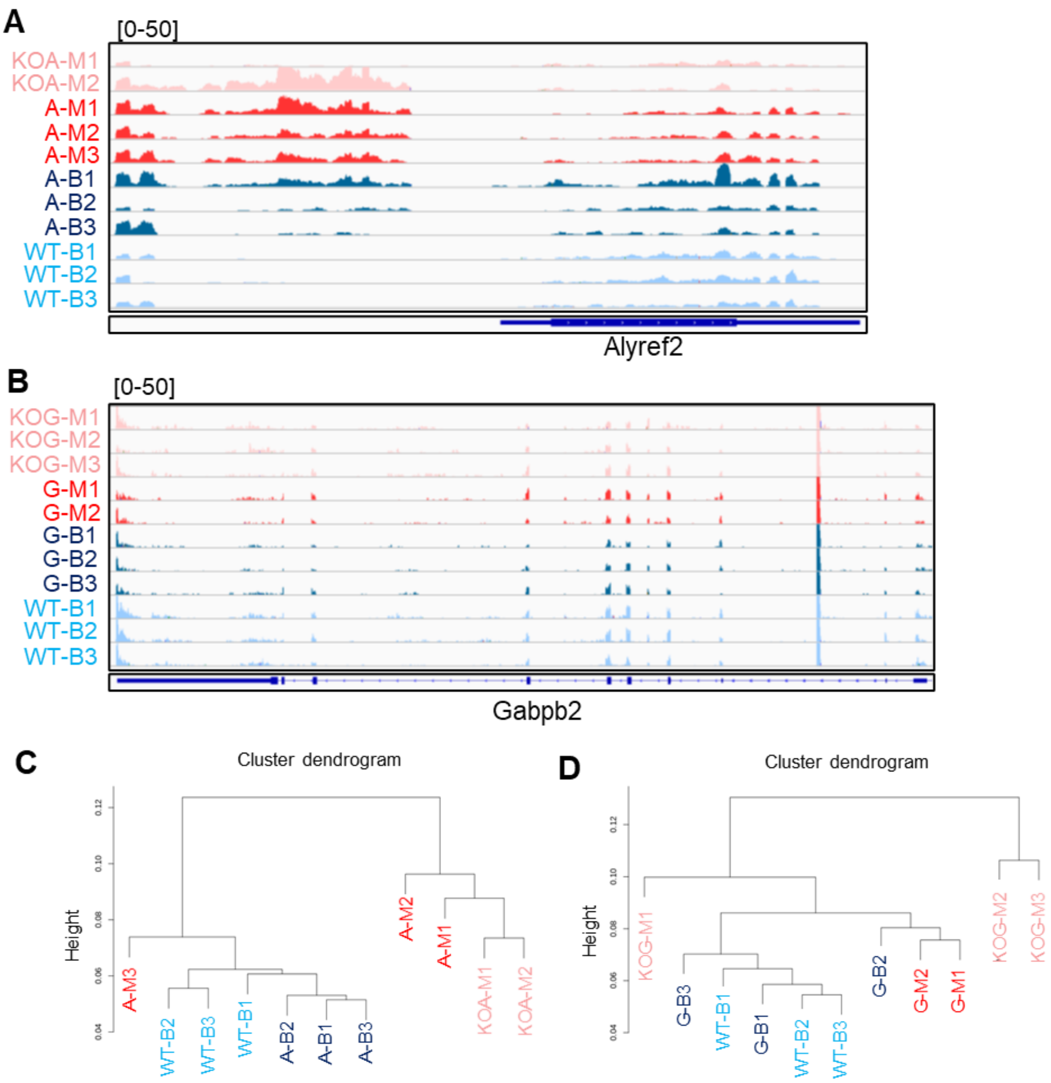


**Figure S5. Single embryo RNA-seq analyses after IVF of *Alyref* and *Gabpb1* heterozygous mice, related to Figure 5**

(A and B) Track images of RNA-seq reads at *Alyref2* (Figure S5A) and *Gabpb2* genes (Figure S5B). Sample names shown here correspond to those in Figures 5B and 5C, respectively.

(C and D) Unsupervised hierarchical clustering analyses of the global gene expression profile among different samples. Sample names shown here correspond to those in Figures 5B and 5C, respectively.

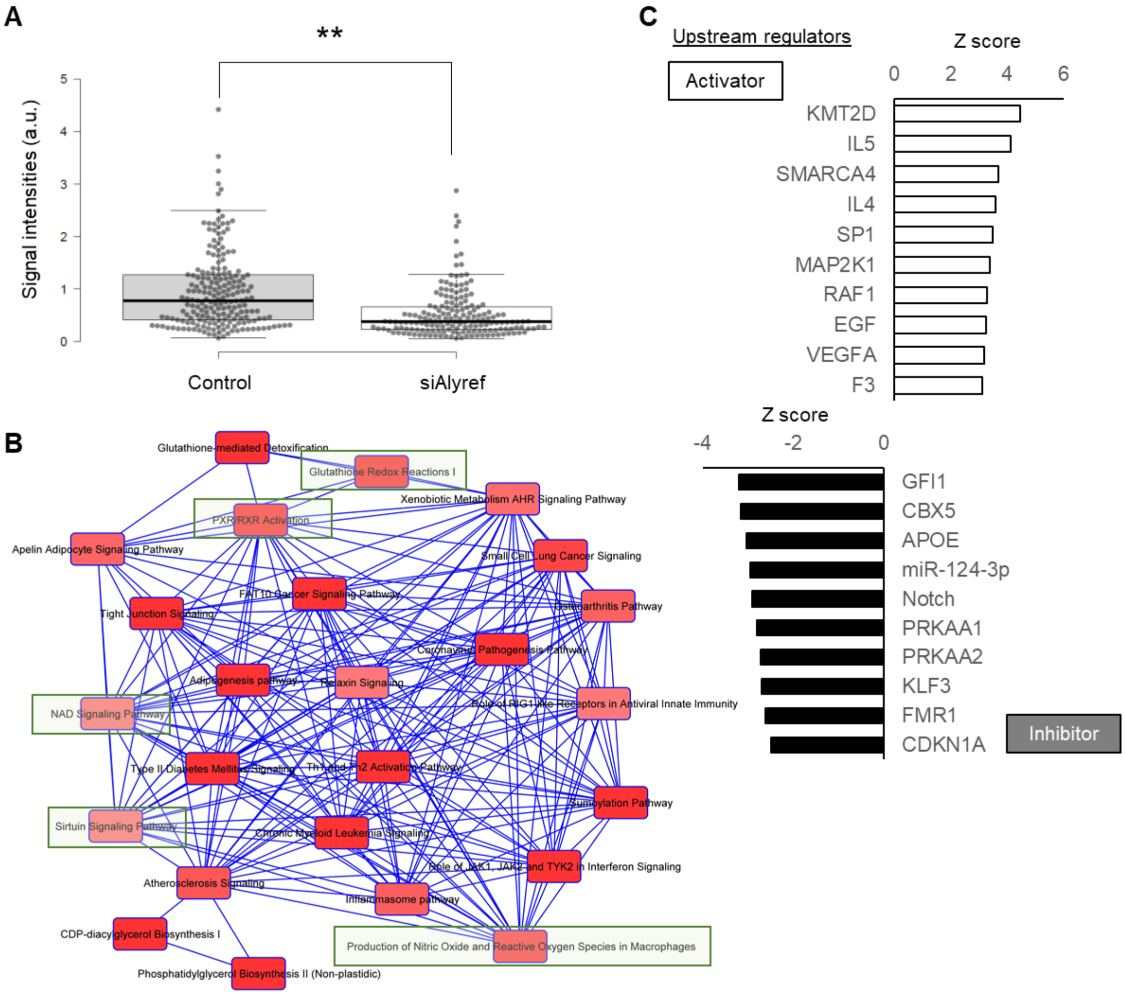


**Figure S6. Misregulated gene expression programs in *Alyref*$^{-/-}$ and *Gabpb1*$^{-/-}$ embryos, related to Figure 6**

(A) Signal intensities of Pou5f1 in embryonic nuclei were compared between siAlyref-injected and control siRNA-injected embryos. Four independent experiments were performed. **p<0.01. p values are based on two-sided T test.

(B) Canonical pathways predicted by IPA using DEGs between *Gabpb1* knockout and wild type (and/or heterozygous) morulae (KOG-M1-3 vs G-M1-2). Closely related terms are connected to each other. Green-highlighted boxes indicate pathways related to anti-oxidation and metabolism as explained in the relevant text.

(C) Predicted upstream regulators responsible for misexpression in Gabpb1$^{-/-}$ embryos were identified by IPA. Activation Z scores are shown.

**Table S1. The list of primers and siRNA**

| Primer name | Sequence (5'-3') | Analysis |
|---|---|---|
| Agpat9 | GGUAAAAUCCGCUAUCGCUGU | siRNA |
| Alyref | GAAACAACUUCCCGACAAAUG | siRNA |
| Bag5 | CACCCACACCGCAUCGAAAUC | siRNA |
| Dr1 | GGGAAAUAAACGAAAGCUAAU | siRNA |
| Gabpb1 | GCACUCCAUACCAACCAGUGG | siRNA |
| Gtf2f2 | CAGAUAAACUGUCAUUGGAAG | siRNA |
| Hipk1 | GCCACUCAAGUACAUAAGACC | siRNA |
| Prmt6 | GAGCACUCUAAUCUAAUAAAA | siRNA |
| Prps1l3 | GCCCAUGAGUAGUACGUUUUC | siRNA |
| Rfpl4b | GCUUUGAUUGCGGUAAAAAAU | siRNA |
| Rnf11 | GGUUUGUAUAACCACAUGACU | siRNA |
| Rps19bp1 | CUAAGUCCGCUCUAGCUGAGU | siRNA |
| Taf9 | GACUAUGAUAAUAUGUAGUCU | siRNA |
| Tle4 | CUGAUGGUCGCACCUUAAUUG | siRNA |
| Zfp939 | GAGUUACUCAUGUAAGAAAAG | siRNA |
| qAlyref_F1 | AGCAAATCCACACAAGTAACAGGA | qPCR |
| qAlyref_R1 | CTAGGCATCACAGGCGAGG | qPCR |
| qGabpb1_F2 | AATTTGCACTCCATACCAACCAG | qPCR |
| qGabpb1_R2 | CGTCACAATGATGGGCTGAC | qPCR |
| qGapdh_F | TCAACGACCCCTTCATTGAC | qPCR |
| qGapdh_R | ATGCAGGGATGATGTTCTGG | qPCR |
| Alyref_F-2 | TGGACATGTCTTTGGACGAC | Genotyping |
| Alyref_R-2 | GACAGTGGGAAGGAGACAGG | Genotyping |
| Gabpb1_Forward-5 | GAAAGGATAACTGAGAATGG | Genotyping |
| Gabpb1_Reverse-5 | CCGTAACAAGATCTGACGCC | Genotyping |
| IF-Alyref-N_Fw | CCCCCTTCACCGCCGACAAAATGGACATGT | Cloning |
| IF-Alyref-N_Rv3 | CCCACCCTTTCATCATTAGCTGGTGTCCATCCTTG CATTGTAAGCATC | Cloning |
| IF-Alyref-vectorF | TGATGAAAGGGTGGGCGCGC | Cloning |
| IF-Alyref-vectorR | GGTGAAGGGGGCGGCCGCGG | Cloning |
| GW_Gabpb1_N_F | CACCTCCCTGGTAGATTTGGGGAAG | Cloning |
| GW_Gabpb1_N_R | CTAAACGGCTTCTTTGTTGG | Cloning |
| GW_Gabpb1_C_F | CACCATGTCCCTGGTAGATTTGGGGAAG | Cloning |
| GW_Gabpb1_C_R | AACGGCTTCTTTGTTGGTCTGG | Cloning |
